# Atlas-Level Single-Cell and Spatial Transcriptomics Data Integration via PRIME

**DOI:** 10.64898/2026.05.20.726698

**Authors:** Xinchao Wu, Xuefei Wang, Jieqiong Wang, Shibiao Wan

**Author notes:** Correspondence: Shibiao Wan.

## Abstract

Single-cell RNA sequencing (scRNA-seq) and spatial transcriptomics (ST) have enabled atlas-scale cellular cartography, with consortium efforts now assembling millions of cells across diverse tissues, donors, and technologies to build comprehensive references for cell identify and disease mechanism, yet the scientific value of these atlases hinges on robust computational integration across heterogeneous data sources. Unlike pairwise batch correction, atlas-level integration must jointly reconcile heterogeneous and often hierarchically nested batch effects across many datasets whose cell-type compositions are highly imbalanced, all while preserving subtle biological variation and remaining computationally tractable at the scale of millions of cells. Existing approaches often prioritize either batch mixing or preservation of local biological structure, and most cannot natively accommodate spatial coordinates. Here we introduce PRIME (Projection-based Robust Integration via Manifold Embedding), an ensemble integration framework that combines random-projection-based consensus anchoring, graph-Laplacian correction, and optional spatial-neighborhood regularization. Across multiple random projections of the expression manifold, PRIME uses consensus voting to keep only cell pairs that repeatedly matched, reducing false anchors caused by projection-specific distortions. For ST, PRIME couples this expression-based anchor graph with a coordinate-derived spatial neighborhood graph in a unified graph-Laplacian objective with closed-form solution, enabling simultaneous cross-batch alignment and local spatial coherence. Based on extensive benchmarking spanning diverse datasets, we show that PRIME consistently outperforms state-of-the-art methods in both batch correction and biological conservation across scRNA-seq and ST integration scenarios and downstream tasks including trajectory inference, spatial-domain preservation, and perturbation-response analysis. Particularly, when integrating a human hematopoiesis benchmark spanning eight donors and approximately 33,000 cells, PRIME preserves biologically coherent developmental trajectories in human hematopoiesis. It also maintains cortical laminar architecture across dorsolateral prefrontal cortex sections in a ST dataset and recovers known drug-target relationships in a perturbation atlas of more than 1 million cells while suppressing batch-associated confounders. Together, these results establish PRIME as a versatile and scalable framework for atlas-level integration of scRNA-seq and ST across diverse biological applications.

## Introduction

Single-cell and spatial transcriptomics now routinely profile millions of cells across tissues, conditions and organisms, enabling systematic comparisons of cellular states and spatial contexts^1,2^. Large reference efforts, including the Human Cell Atlas^1^ and Tabula Sapiens^3^ serve both as descriptive catalogs of cellular diversity and as frameworks for comparative and mechanistic analyses across studies^1,3,4^. Building such atlases, however, requires the integration of datasets generated with different dissociation protocol, library preparation methods, sequencing chemistries and depths, introducing technical variations that can approach or exceed the biological differences of interest^5,6^. The challenge of atlas-level integration is therefore not simply to mix datasets, but to selectively remove technical variation while retaining biologically meaningful structure, including rare transitional populations, developmental trajectories, perturbation-induced expression programmes and tissue organization. This balance has bilateral benefits: undercorrection fragments equivalent cell states by study of origin, whereas overcorrection can spuriously merge distinct cell populations or attenuates context-dependent signals. Moreover, neither failure mode can be diagnosed reliably from batch-mixing metrics alone^7^.

These limitations have become increasingly consequential as downstream analyses have shifted from cluster annotation toward problems that depend on local manifold structure, temporal trajectory inference, and spatial context. In developmental systems, batch-induced distortions in neighborhood relationships can misdirect trajectory inference algorithms such as Monocle^8,9^ or PAGA^10^, producing spurious branch points or inverting the ordering of transitional states^11^. In spatial transcriptomics, transcriptionally similar cells located in distinct anatomical regions may represent functionally distinct states, such that methods that ignore spatial coordinates risk merging them inappropriately^12–14^. In large-scale chemical or genetic perturbation screens, where hundreds of conditions are profiled across multiple batches, over-correction can suppress the perturbation signatures one aims to characterize, particularly when matched control are limited^15–17^. Together, these settings define complementary and stringent requirements for integration that extend beyond standard benchmarks of batch removal. Yet the extent to which current methods preserve these properties after correction has not been systematically evaluated, and no general framework is designed to address all three settings while accommodating both dissociated and spatial data.

A broad range of integration methods for integrating scRNA-seq or spatial transcriptomics has been developed, yet these approaches differ substantially in algorithmic assumptions and scalability. For dissociated single-cell data, mutual nearest-neighbor (MNN) matching (e.g. fastMNN^18^) provide a conceptually simple and computationally efficient strategy, but their performance can be sensitive to distortions in the embedding space used for matching. Anchor-based alignment method, including CCA^19^, are effective for correcting substantial batch effects, although noisy or imbalanced anchor sets may lead to overcorrection and obscure fine-grained cellular heterogeneity. Iterative linear correction frameworks such as Harmony^20^ scale well to large datasets and promote effective batch mixing, but may attenuate rate transitional or subtly distinct cell states. Graph-based approaches, like BBKNN^21^, are highly efficient, yet primarily modify neighborhood structure rather than producing an explicitly corrected embedding. And deep generative models including scVI^22^, offer flexible nonlinear integration, but demand substantial computational resources, careful model specification, and can be difficult to diagnose when performance degrades at atlas scale. For spatial transcriptomics, an emerging set of methods explicitly leverages spatial coordinates. Optimal-transport-based alignment such as PASTE^23^ are well suited to aligning consecutive tissue sections, but typically rely on anatomical correspondence across slices. While graph-based representation learning methods, including GraphST^24^ and STAligner^25^, can integrate multiple spatial samples without strict pre-alignment, but their performance may depend on training stability and graph construction. Thus, despite substantial progress, most single-cell integration methods do not explicitly model spatial context, whereas spatially-aware methods are not generally designed for atlas-scale integration. This separation leaves an important methodological gap: a unified and principled framework capable of operating across both dissociated and spatial transcriptomic regimes.

Here we present PRIME (Projection-based Robust Integration via Manifold Embedding), a unified graph-Laplacian framework that performs robust transcriptomic integration with a single mathematically self-consistent formulation for two regimes: atlas-level scRNA-seq integration across millions of dissociated cells, and multi-slice integration of spatial transcriptomics data. The expression-based anchor graph is built by consensus voting across an ensemble of independent random projections, in which only correspondences supported by most projections survive, suppressing the projection-specific distortions that destabilize single-embedding methods. The spatial neighborhood graph, constructed from tissue coordinates when available, captures within-slice geometric proximity. The two graphs are reconciled in a closed-form, convex graph-Laplacian objective that simultaneously enforces cross-batch alignment and within-tissue spatial smoothness; when only dissociated data are provided, the spatial term reduces to zero so that the same mathematical formulation seamlessly scales from atlas-level scRNA-seq to multi-slice spatial integration. We first demonstrate on a controlled mixture cell-line dataset and a large-scale lung cancer atlas that PRIME outperforms existing methods across both batch correction and biological conservation metrics. We then show that PRIME retains biologically coherent developmental trajectories in human hematopoiesis, which is a stringent test that existing benchmark metrics do not systematically assess. In the ST data integration, PRIME preserves cortical laminar architecture when integrating multiple DLPFC tissue sections. Finally, applying PRIME to a large-scale single-cell perturbation atlas comprising over 1 million cells, we show that it recovers known drug–target relationships while suppressing batch-associated confounders. Together, these analyses demonstrate that stabilizing local neighborhood structure during batch correction is essential for preserving the biological resolution required for developmental, spatial, and perturbational downstream analyses.

## Results

### PRIME enabled robust integration of single-cell and spatial transcriptomics through ensemble random projection anchors and graph-Laplacian smoothing

We developed PRIME (Projection-based Robust Integration via Manifold Embedding), a unified framework that integrates single-cell and spatial transcriptomics datasets across batches by combining two complementary signals: cross-batch mutual-nearest-neighbor (MNN)^18^ anchors aggregated over an ensemble of random projections^26^, and within-tissue spatial proximity encoded by a coordinate-based neighborhood graph (**Fig. 1**). The two signals are reconciled in a single graph-Laplacian^27^ objective whose closed-form solution yields a batch-corrected expression matrix that removes technical variation while preserving both cell-type heterogeneity and tissue-level spatial continuity. PRIME takes normalized expression matrices and batch labels from two or more datasets as input. For spatial transcriptomics data, two-dimensional spot or cell coordinates are additionally supplied to construct a spatial neighborhood graph^12^. The output is an integrated embedding that can be used consistently for clustering, trajectory inference, spatial-domain identification, perturbation analysis, and benchmark evaluation.

**Figure 1.**
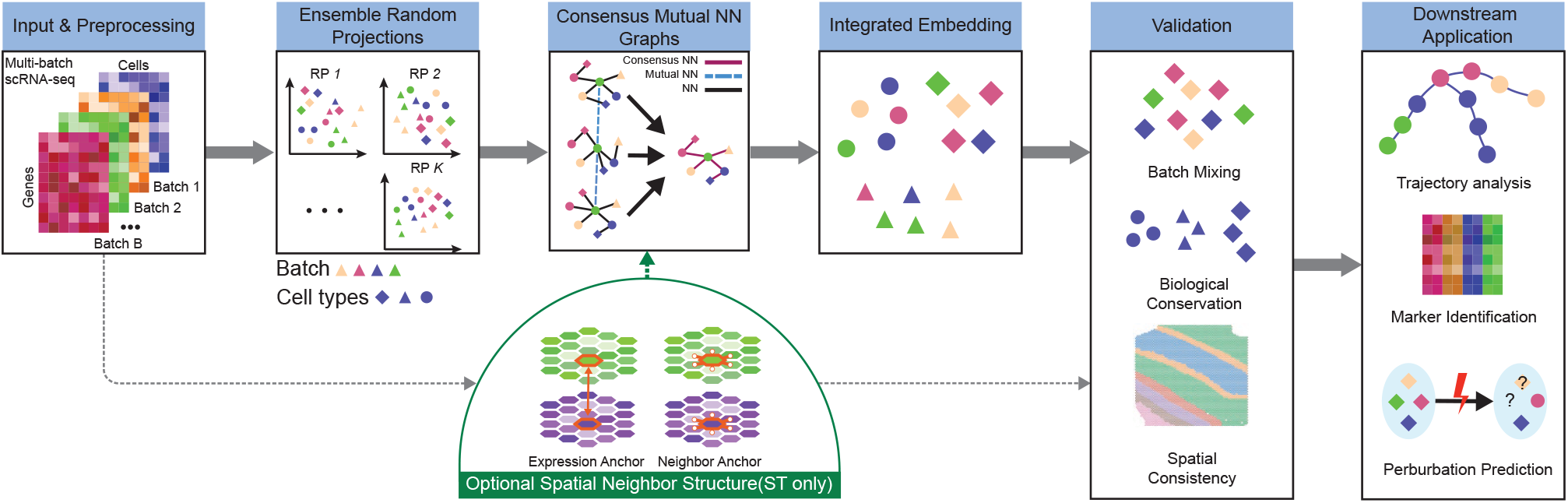
Overview of the PRIME framework for single-cell and spatial transcriptomics integration. PRIME takes normalized multi-batch expression matrices as input and constructs an ensemble of random-projection views. Within each projected space, mutual nearest-neighbor anchors are identified between batches, and anchors supported across multiple projections are retained as consensus correspondences. The consensus expression graph is used to generate a batch-corrected embedding that aligns shared biological states across batches while preserving local cell-state structure. For spatial transcriptomics data, PRIME additionally incorporates two-dimensional tissue coordinates by constructing a spatial neighborhood graph and combining it with the expression-based anchor graph through graph-Laplacian regularization. The resulting integrated representation supports downstream clustering, trajectory inference, spatial-domain analysis, perturbation analysis, and quantitative benchmarking.

The PRIME workflow comprises five steps (**Fig. 1**). (1) Random projection ensemble: the input expression matrix is independently projected *K* times into lower-dimensional spaces using Gaussian random matrices^26,28^, producing an ensemble of views of the data. Previously, we have extensively applied random projection theory in other areas like scRNA-seq clustering^29^. (2) Per-projection MNN identification: within each projected view, mutual nearest neighbors^18^ are identified between batch pairs, yielding *K* candidate anchor sets. (3) Consensus anchor selection: candidate pairs are aggregated across projections via majority voting, retaining only those supported by a fraction *τ* of views to form an expression-based anchor graph. (4) Spatial graph construction: for ST data, an additional neighborhood graph is built by connecting each cell or spot to its *k*_*s*_ nearest spatial neighbors with Gaussian-weighted edges. (5) Laplacian-regularized correction: the expression-based and spatial graphs are jointly embedded into a graph-Laplacian objective whose minimization yields the corrected embedding. Intuitively, PRIME treats each random projection as an independent view of the data’s local geometry and trusts only those anchors with consensus votes, while the spatial graph acts as a geometric prior that keeps physically adjacent cells close in expression space even after correction, preventing anchor-based alignment from erasing the smooth gradients that define tissue architecture.

Specifically, PRIME rests on three principal ingredients: ensemble random projection, consensus-based anchor voting, and graph-Laplacian regularization. Let **X** ∈ ℝ^*n*×*d*^ denote the normalized expression matrix over *n* cells and *d* genes. PRIME first generates *K* projected embeddings

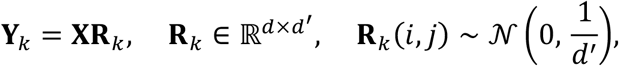

where **R**_*k*_ is the *k*-th Gaussian random matrix, **Y**_*k*_ is the resulting projected embedding for the *k*-th view, and *d*′ is the dimension of the lower-dimensional projection space (default *d*′ = 128, and *d*′ ≪ *d*). The Johnson–Lindenstrauss lemma^26^ guarantees that pairwise distances are approximately preserved under such projections with high probability, so that local neighborhood structure is retained across all *K* views. Within each view **Y**_*k*_, mutual nearest neighbors between two batches of single cell data sets 𝔄 and 𝔅 are defined by the symmetric condition.

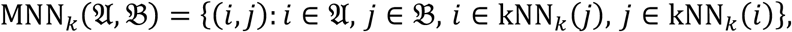

which suppresses spurious anchors arising from sampling imbalance. Aggregating votes across projections, 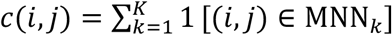, and retaining pairs with *c*(*i, j*) ≥ *τK* (default *τ* = 0.5) yields the expression-based anchor graph *G*_*E*_ = (𝔙, *E*_*E*_, W_*E*_) with edge weights W_*E*_(*i, j*) = *c*(*i, j*)/*K*. Here, the scalar count *c*(*i, j*) records the number of projections in which the cell pair (*i, j*) appears as a mutual nearest neighbor; the vertex set 𝔙 contains all cells across batches, the edge set *E*_*E*_ collects the retained consensus anchor pairs, and the weight matrix W_*E*_ stores the support-fraction edge weights.

For batches with spatial coordinates {*p*_*i*_ ∈ ℝ^2^} (where the vector *p*_*i*_ denotes the two-dimensional spatial location of cell *i*), PRIME additionally constructs a spatial graph *G*_*S*_ = (𝔙, *E*_*S*_, W_*S*_) in which each cell is linked to its *k*_*s*_ nearest spatial neighbors with Gaussian-weighted edges,

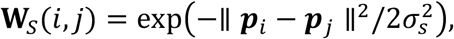

where *σ*_*s*_ is set adaptively to the median distance among the *k*_*s*_ neighbors. The two graphs are then combined into a single objective whose minimization produces the corrected embedding 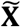:

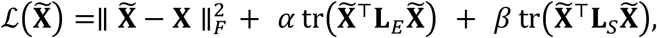

where **L**_*E*_ and **L**_*S*_ are the normalized Laplacians of the expression and spatial graphs, respectively, and the trace form tr 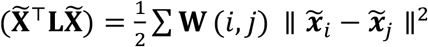penalizes disagreement between graph-connected cells. **W**(*i, j*) is the desired weighted matrix, and 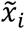 is the corrected embedding of *i*-s cell. The objective is convex and quadratic, admitting the closed-form solution:

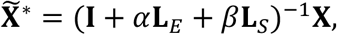

which we compute efficiently using sparse conjugate-gradient solvers. The three terms have distinct biological roles: data fidelity anchors the corrected profiles to the measurements, cross-batch alignment pulls consensus anchors from different batches toward each other, and spatial smoothness preserves expression continuity between physically neighboring cells, preventing the correction from disrupting tissue-layer transitions or tumor–stromal boundaries. When spatial coordinates are unavailable, setting *β* = 0 reduces the framework to pure expression-based integration without any change in implementation.

PRIME achieves significant improvements over other methods. Classical MNN-based methods (e.g., fastMNN^18^ identify anchors in a single embedding space and are therefore sensitive to projection-specific distortions, while recent spatial integration methods (e.g., PASTE^30^, GraphST^24^, STAligner^25^) rely primarily on spatial coordinates and cannot gracefully handle expression batches. PRIME unifies both regimes by treating expression-based consensus anchoring and coordinate-based spatial regularization as two graphs inside a single Laplacian objective, yielding a method that is applicable to both scRNA-seq and ST data integration. The framework exposes only five interpretable hyperparameters (*K, τ, k*_*s*_, *α, β*), with a convex closed-form solution, making it simultaneously lightweight and theoretically well-grounded. A complete derivation is provided in the Methods section.

### PRIME outperformed state-of-the-art methods across simulated and real single-cell datasets

To systematically evaluate PRIME, we first evaluated whether both the ensemble random projection and the consensus voting over anchors contribute meaningfully to integration quality. We used a classical 10x Genomics mixture-cell-line dataset comprising approximately 3,000 cells, in which Jurkat and HEK293T cells were profiled both as two pure, single-cell-line batches and as a 1:1 mixture batch^31^ (**Supplement Fig. 1**). This setup provides an unambiguous ground truth for cross-batch correspondence. On this dataset, we compared three configurations: (i) uncorrected PCA on the concatenated expression matrix, (ii) MNN anchoring in a single random projection (single RP), and (iii) the full PRIME framework with consensus voting. Uncorrected PCA failed to align the three batches, producing four disjoint clusters (**Suppl Fig. 1A**). Single RP recovered partial mixing but introduced visible anchor errors between Jurkat and 293T cells, reflecting the sensitivity of any one random projection to local geometric distortions (**Suppl Fig. 1B-C**). Only the full PRIME framework achieved near-perfect batch mixing while preserving the two-cluster cell-line structure, confirming that the ensemble of projections is essential to preserve the biological structure (**Suppl Fig. 1D**).

To further evaluate the performance of PRIME in real-world large-scale scRNA-seq data integration, we next benchmarked with 10 state-of-the-art methods on a substantially more challenging dataset: a human lung cancer scRNA-seq compendium comprising 892,296 cells from 44 cell types from 12 studies, encompassing tumor, stromal, and immune cell populations^32^. We benchmarked PRIME against ten widely used single-cell integration methods spanning four algorithmic families: classical MNN-based (MNN, FastMNN, Scanorama^33^), linear mixed-effects models (Harmony^20^, ComBat^34^, CCA^19^) deep-learning-based (scVI^22^), graph-based (BBKNN^21^), and matrix-factorization-based (jPCA and RPCA^35^) as shown in **Fig. 2A-B**. To ensure a fair and reproducible comparison, we evaluated all methods using scIB^5^. The framework jointly quantifies two competing objectives: batch mixing (how well technical variation is removed, measured by iLISI, kBET, batch ASW, and graph connectivity) and biological conservation (how well cell-type structure is preserved, measured by cLISI, NMI, ARI, and isolated-label scores). scIB further aggregates these complementary axes into a single composite score, enabling a principled ranking across methods. Compared to the other methods, PRIME ranked first or tied for first on total aggregated scores, outperforming the second-best method by 2% on the overall composite score (**Fig. 2C**). Importantly, PRIME achieved this leading performance while simultaneously attaining the highest batch correction scores (iLISI = 0.31, kBET = 0.34) and among the highest biology preservation scores (cLISI = 1.00, NMI = 0.76, ARI=0.65), whereas most competitors traded one objective against the other. For example, ComBat achieved strong batch mixing at the expense of biological conservation, while FastMNN showed the opposite pattern (**Fig. 2C**). These results demonstrate that the PRIME generalizes from controlled two-cluster benchmarks to heterogeneous multi-donor cancer atlases, and that PRIME resolves the batch correction versus biology preservation trade-off more effectively than existing methods.

**Figure 2.**
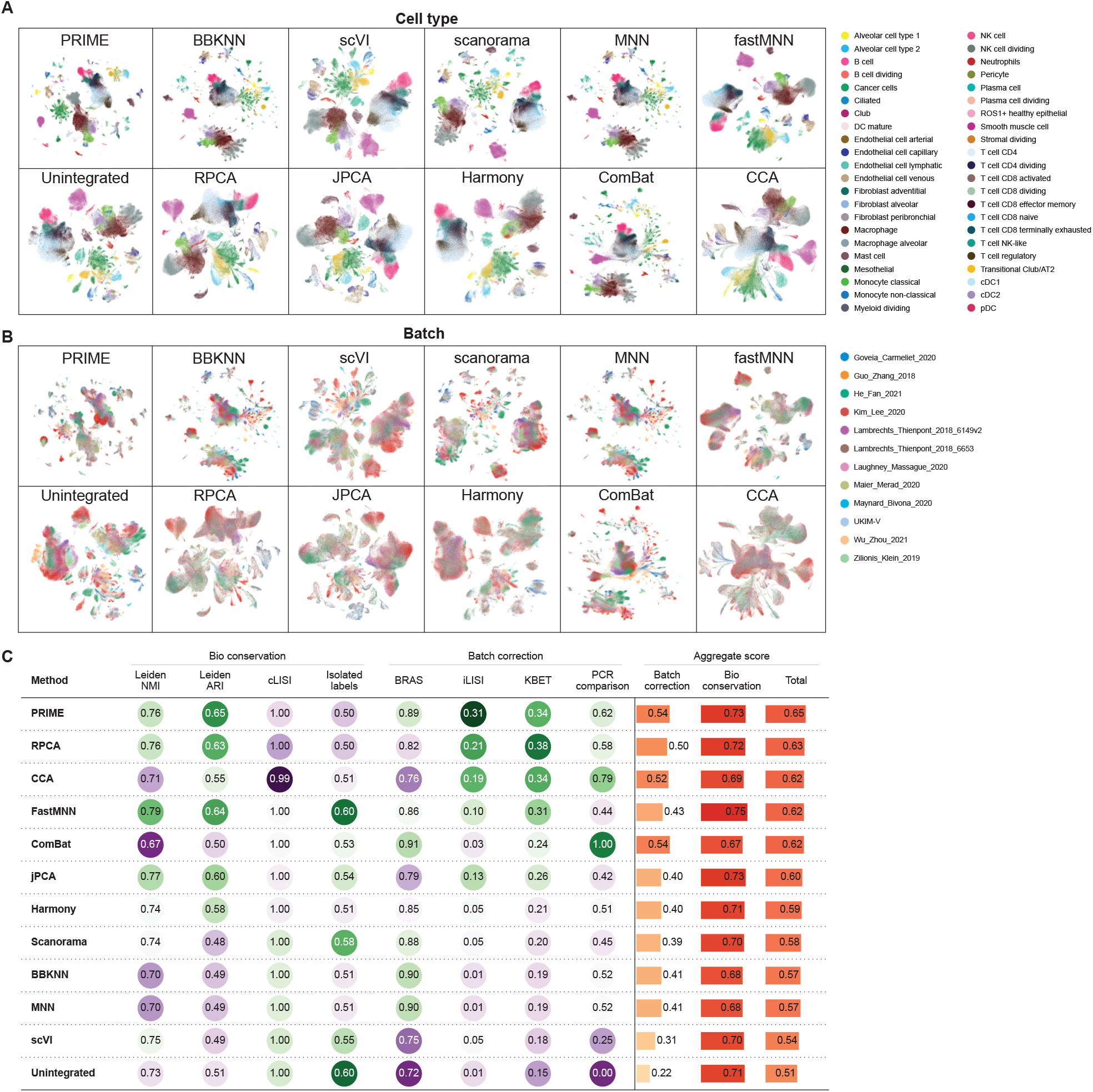
PRIME achieved strong batch correction while preserving biological structure across single-cell benchmarks. Overview of the real-world lung cancer integration benchmark across donors, studies, and cell populations. The atlas contains 892,296 cells from 12 datasets. **A-B**, Integrated embeddings generated from different methods, cells colored by cell types and sources, respectively. **C**, Quantitative scIB benchmark table comparing PRIME with ten single-cell integration methods. Metrics are grouped into biological conservation and batch correction categories, scaled from 0 to 1 (the higher the better), and ordered by total aggregate score. Aggregated benchmark scores summarize the balance between batch mixing and biological conservation across methods.

### PRIME revealed underlying lineage after batch correction

To evaluate whether PRIME preserves developmental and differentiation trajectories, we applied it to a human hematopoiesis dataset comprising 33,506 cells from 8 donors profiled by 10X Genomics (**Supplement Fig. 2a-b**). The dataset includes a well-characterized developmental hierarchy spanning hematopoietic stem cells (HPSCs), multipotent progenitors (MPPs), and lineage-committed precursors (e.g., erythroid, myeloid, lymphoid). We first benchmarked PRIME against the same panel of methods as in the previous section, using scIB to quantify batch mixing and biological conservation. PRIME achieved the highest overall aggregate scores, same as the second-best method but outperforming the third-best by 1% (**Fig. 3A**). Even though the superior performance of PRIME cannot directly reflect into leading in metrics, the underlying trajectory structure was revealed clearly in the PRIME generated integrated embedding. To assess trajectory preservation, we applied Monocle 3^36^ to the integrated embeddings produced by PRIME and compared the inferred trajectories to known hematopoietic lineages (**Fig 3B-C**). Specifically, there are two well-established branches in hematopoiesis^37,38^: the erythro-myeloid branch, which gives rise to erythrocytes and myeloid cells, and the lymphoid branch, which gives rise to B cells, T cells, and NK cells. Here, we focused on the erythroid–myeloid continuum, where progenitor-to-lineage transitions are gradual and therefore particularly sensitive to overcorrection. We used HSPC/MPP populations as the trajectory root and evaluated whether cells progressed toward erythroid and myeloid/monocyte termini with expected marker-gene dynamics (**Fig. 3C**). In contrast, several other methods, including Harmony and fastMNN, produced embeddings in which the continuous structure was disrupted, and the gap between lineages was artificially closed, consistent with overcorrection (**Supplement Fig. 2A-B**). These results demonstrate that PRIME’s corrected embedding retains the expected developmental structure without requiring additional tuning or post hoc trajectory-specific adjustments.

**Figure 3.**
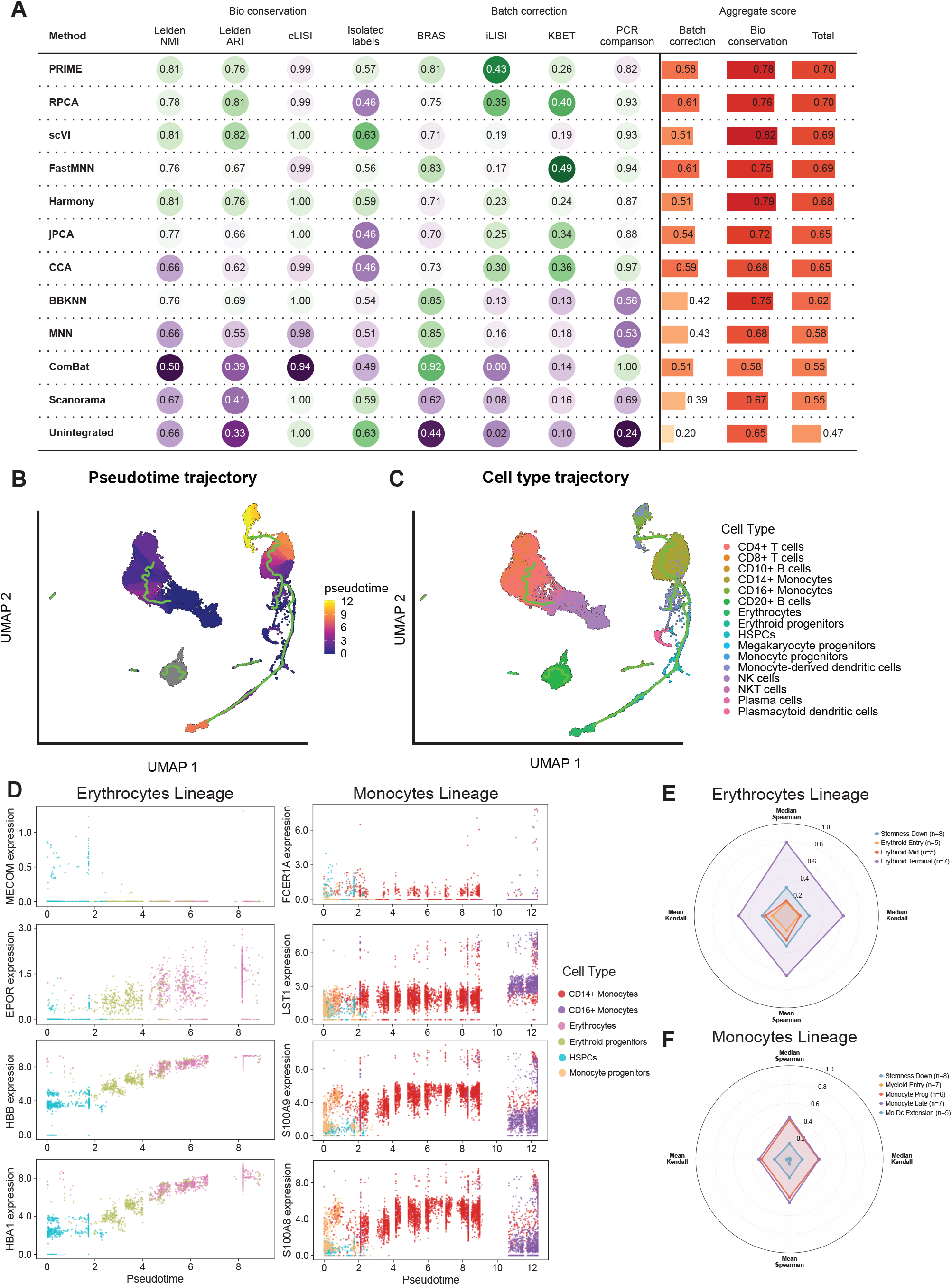
PRIME preserved hematopoietic lineage topology and lineage-specific gene dynamics after batch correction. **A**, Benchmarking summary comparing PRIME with alternative integration methods on batch correction and biological conservation metrics on human immune cell atlas including 33,506 cells from 8 donors. **B-C**, Monocle 3 trajectory inferred from the PRIME-corrected embedding, showing a preserved erythroid–myeloid bifurcation with HSPC/MPP cells near the root and lineage-committed cells at the termini. **D**, Expression or module-score trends of known hematopoietic markers along pseudotime, including decreasing HSPC-associated programs and branch-specific enrichment of erythroid and myeloid/monocyte markers. **E-F**, Quantitative trajectory-conservation summary based on marker-gene dynamics and pseudotime-associated gene correlation.

Furthermore, we evaluated the conservation of known lineage-specific gene programs after integration, and applied metrics including gene-gene correlation preservation and lineage-specific marker gene expression. The erythroid branch was evaluated using canonical erythroid markers such as GYPA, HBA1, HBA2, HBB, ALAS2, and SLC4A1, whereas the myeloid/monocyte branch was evaluated using markers such as LYZ, FCN1, S100A8, S100A9, LST1, and CTSS^38^ (**Fig. 3D**). And the high score of conservation of pseudo-time gene-gene correlation (**Fig. 3E-F**) indicates that the relative ordering of gene expression along the trajectory is also preserved, which is critical for downstream analyses such as gene regulatory network inference. Together, these results demonstrate that PRIME not only achieves strong batch correction and cell-type preservation, but also retains the continuous developmental structure and lineage-specific gene programs that are essential for understanding biological processes.

### PRIME integrated spatial transcriptomics slices while preserving tissue architecture

Joint analysis of multiple slices from matched samples across donors or conditions is essential for constructing spatial atlases and comparing tissue architecture across biological contexts^39^. But it is complicated by slice-level batch effects that can distort or obscure the underlying tissue structure. Unlike scRNA-seq integration, ST-to-ST integration must simultaneously remove these batch effects and preserve the spatial coherence within each slice, so that tissue domains remain well-defined in the corrected embedding^30^.

To systematically evaluate the spatial integration capabilities of PRIME, we applied it to a human dorsolateral prefrontal cortex (DLPFC) dataset comprising 12 tissue sections profiled by 10x Visium, each containing approximately 4,000 spots^39^. The DLPFC is a well-studied brain region with a well-characterized six-layer laminar architecture and manually annotated cortical layer labels, making it an ideal test case for spatial integration because ground-truth tissue domains can be used to rigorously assess whether integration preserves biologically meaningful structure. We benchmarked PRIME against four other state-of-the-art spatial integration methods, including GraphST^24^, PRECAST^40^, STAligner^25^, and PASTE^30^. In the joint UMAP embedding, spots integrated by PRIME co-localized by cortical layer rather than by slice of origin, with spots from each annotated layer forming compact, well-mixed clusters that bridged all 8 sections (**Fig. 4A-B**). In contrast, GraphST exhibited totally mixture of cortical layers, indicating incomplete cross-slice alignment, and PRECAST ignored the spatial coherence across the cortical layers. PSATE was designed to align the spatial structures of the cross-slices data, but the integrated embedding exhibited no spatial structure. To assess integration quality, we evaluated three complementary aspects using metrics drawn from the scIB framework, and our purposed cross-layer continuity (XLC) metric. We encoded the DLPFC layer annotations as an ordinal variable, with Layer1, Layer2, …, Layer6 assigned ranks 1, 2, …, 6, and white matter assigned as rank 7, reflecting its anatomical proximity to Layer6. Quantitatively, PRIME outperformed STAligner and other methods in the total score, aggregating batch correction, bio conservation and spatial continuity (**Fig. 4C**). These results demonstrated that the Laplacian-based spatial smoothness term successfully prevents batch correction from disrupting within-slice tissue architecture.

**Figure 4.**
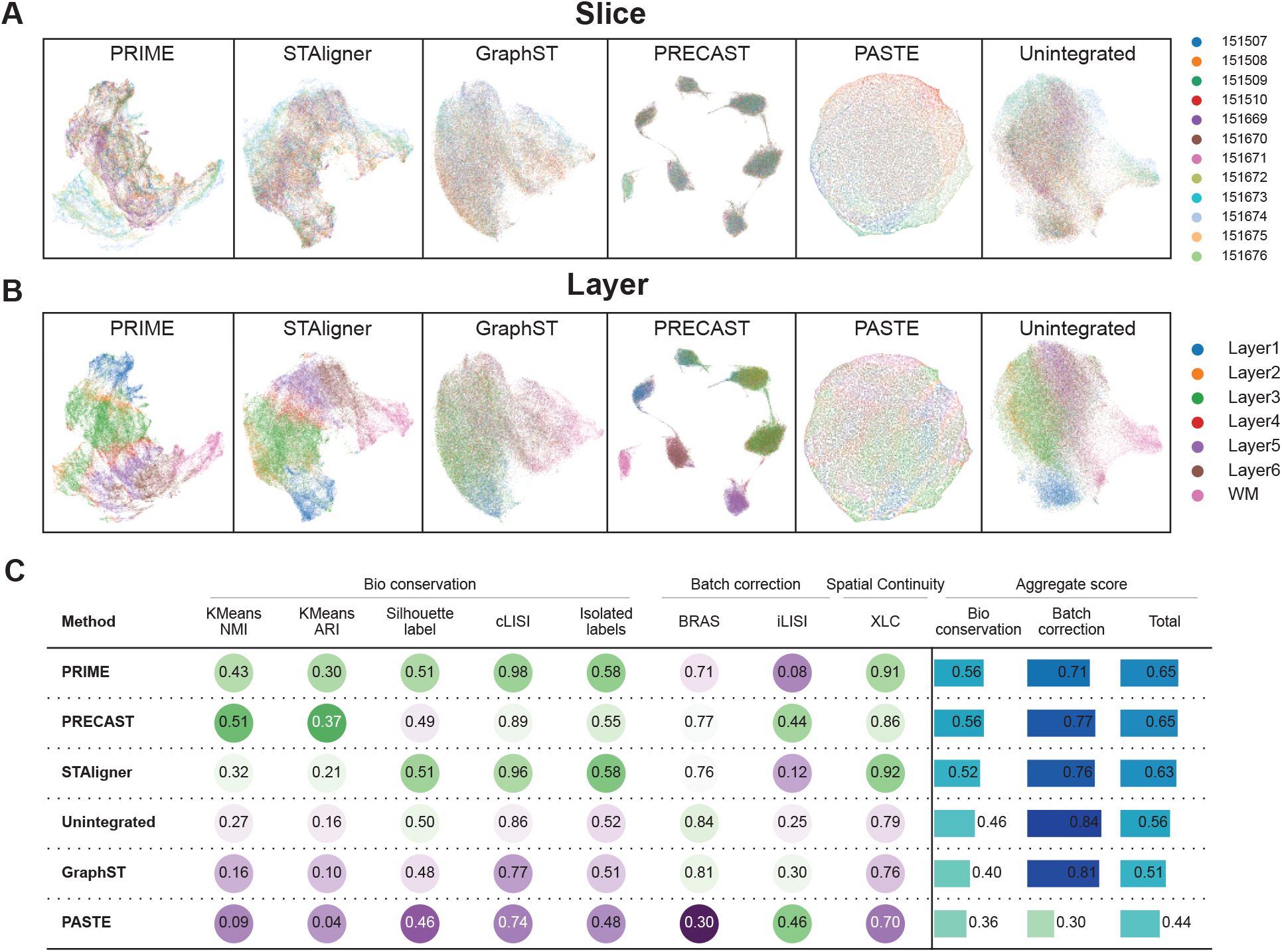
PRIME integrates multiple DLPFC spatial transcriptomics sections while preserving cortical-layer architecture. **A**-**B**, UMAP embedding of PRIME-integrated and other methods, spots colored by the slice identity and layer annotation of the DLPFC data, respectively. The dataset includes 47,681 cells from 12 slices. **C**, Benchmark comparison of PRIME with other methods, using Bio conservation, Batch correction and Spatial Continuity metrics, ordered by Total aggregated score.

### PRIME facilitated drug target discovery from large-scale perturbation dataset

To further demonstrate the superior performance of PRIME for single cell perturbation response effects, we applied it to the Tahoe-100M dataset^17^, a large-scale perturbation screen comprising more than 100 million single cells, spanning 50 cell lines and 1000 more drug treatments. The dataset was designed to systematically evaluate the effects of a wide range of perturbations on diverse cellular contexts, but its scale and complexity pose substantial challenges for integration and analysis (**Fig. 5a**). The batch effects between perturbed and control group were mixed with the perturbation signal, resulting a confounding technical noise associated with true perturbation effects (**Fig. 5b**). The energy-distance (E-distance) was leveraged to quantify the perturbation effect size, in which a larger E-distance indicates stronger distinction from control group (**Fig. 5c**). We selected to apply PRIME for the lung cancer cell lines, which include 424,498 cells across 30 drugs and dimethyl sulfoxide (DMSO) controls^17^, and evaluated whether PRIME could effectively integrate the data across batches and reveal meaningful biological insights. After applying PRIME to the lung cancer subset, we first evaluated integration quality using perturb separation E-distance and batch residual E-distance, against five other methods, including Harmony, ComBat, CCA, MNN, and scanorama. PRIME achieved the perturb separation E-distance while low batch residual E-distance (**Fig. 5d**), suggesting it mitigate the batch effects while preserve the perturbation signals after correction.

**Figure 5.**
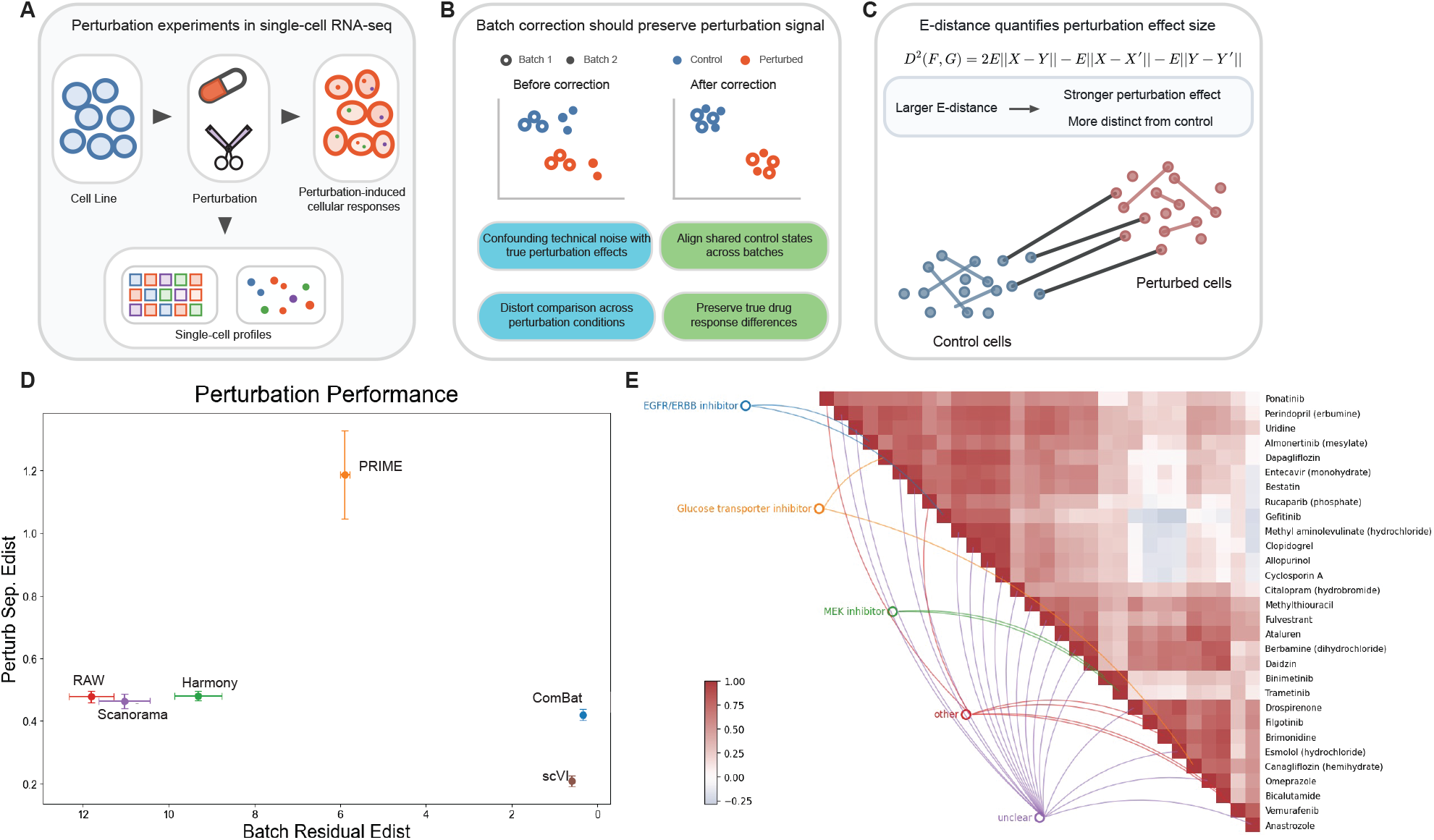
PRIME enables mechanism-of-action analysis in large-scale single-cell perturbation data. **A**, Workflow for single-cell perturbation experiments, perturbed cells are sequenced to generate perturbed profiles alongside DMSO controls. The lung cancer perturbation cell line contains 424,498 cells from 12 samples. **B**, the batch effects in perturbation analysis is confounding factor to perturb effect identification, and a ideal batch effect correction method should preserve drug response signal while mitigate batch effects. **C**, Energy-distance (E-distance) is designed to measure the batch effect and perturb effect in perturbation analysis, a stronger E-distance indicates more distinction from control group. **D**, Benchmark performance of PRIME and competing methods on the Tahoe-100M lung-cancer perturbation subset, evaluated using perturb separation E-distance and batch residual E-distance. **E**. Correlation heatmap summarizing relationships between perturbation-induced transcriptional signatures of different drug perturbation. Strong correlations recover expected drug relationships and nominate potential mechanisms for compounds with unclear or incompletely characterized targets.

After that, to evaluate the utility of PRIME for drug target discovery, we performed differential expression analysis between drug-treated and control cells within the lung cancer-associated cell lines, using the PRIME-corrected embedding to define the differential expression contrasts. Following standard protocols, we identified differentially expressed genes (DEGs) for each drug treatment compared to DMSO controls and then performed correlation analysis between the DEG profiles of each drug. According to the correlation analysis, it revealed several known drug-target relationships (**Fig. 5e**), such as the correlation between the DEG profile of two MEK inhibitor, confirming that PRIME’s integrated embedding supports meaningful biological contrasts. Moreover, there are also many drugs whose target pathways are unknown or poorly characterized, and the correlation analysis revealed strong associations between their DEG profiles and specific pathway signatures, suggesting potential mechanisms of action that warrant further experimental validation. These results demonstrate that PRIME not only achieves strong integration performance on large-scale perturbation datasets, but also enables downstream analyses such as drug target discovery by providing a corrected embedding that supports meaningful biological contrasts.

## Discussion

Constructing coherent cell atlases from heterogeneous single-cell and spatial transcriptomics studies requires integration methods that simultaneously remove technical variation across batches and preserve the fine-grained biological structure required for trajectory, spatial, and perturbational analyses^5,6^. Existing methods struggle with this balance, because aggressive batch alignment distorts the local neighborhood relationships on which downstream inference depends. This failure has concrete biological consequences: overcorrection that spuriously merges distinct cell states can misdirect pseudotime ordering, invert trajectory branches, or eliminate the transcriptional contrasts that perturbation screens are designed to detect. Undercorrection that preserves batch-induced fragmentation prevents the assembly of coherent cross-study references and obscures shared cell states. PRIME can resolve this trade-off by stabilizing cross-batch neighborhood structure, preserving local manifold geometry that trajectory inference, spatial domain identification, and perturbation analysis all rely on. Across diverse batch effect correction settings, PRIME outperformed state-of-the-art methods either on the benchmarking metrics or underlying structure revealed by downstream analysis.

The conceptual basis of PRIME is that any single embedding provides only one view of cell-cell geometry, and view-specific distortions translate directly into cross-batch anchor misalignment. Because independent random projections distort different parts of the manifold, aggregating cross-batch anchors across *K* projections through consensus voting operates as safeguard. An anchor will be selected only if it is stable across multiple independent views, which suppresses projection-specific false positives while retaining biologically robust correspondences. The spatial extension applies Graph Laplacian Regularization, in which the expression graph L_*E*_ enforces cross-batch alignment through consensus anchors, while the spatial graph L_*S*_ enforces the complementary constraint of intra-tissue smoothness. The probabilistic argument underlying consensus voting follows the logic of Condorcet’s jury theorem: if each projection independently has greater than chance probability of identifying a true anchor, then the majority vote over many projections converges on the correct answer with probability approaching one as ensemble size grows. In practice, this means that the false-anchor rate decreases geometrically with the number of projections, explaining why the ensemble outperforms any single projection even when each individual view is only moderately accurate. The spatial extension reformulates the integration as a regularized least-squares problem: the expression-based anchor graph provides the cross-batch alignment signal, while the spatial graph acts as a geometric prior that penalizes corrections which would displace physically neighboring cells apart in expression space. Because both graphs enter the objective as convex quadratic terms, the combined problem admits an efficient closed-form solution via sparse conjugate-gradient solvers, making PRIME computationally tractable for large datasets while remaining principled in its treatment of the spatial-expression trade-off. This design prevents cross-slice correction from disrupting the within-slice laminar organization that defines tissue architecture.

This design yields concrete biological dividends that aggregate benchmark scores do not capture and that existing benchmarks have not systematically evaluated. In trajectory analysis of human hematopoiesis, the continuous erythro-myeloid bifurcation remained resolvable in the corrected embedding, with HSPCs clearly positioned at the root of the trajectory and lineage-committed precursors at the termini. By contrast, other methods produced embeddings in which the gap between committed lineages was artificially closed, leading to misdirected pseudotime inference and downstream gene-regulatory-network analyses. Lineage-specific marker programs (*GYPA* and *HBB* for erythroid, *CLEC4C* and *CLECSA* for myeloid) retained their expected expression gradients along the preserved trajectory, with pseudotime gene–gene correlation conserved. This preservation of pseudotime-correlated gene dynamics is critical because trajectory models are frequently used to infer transcription factor hierarchies and gene regulatory networks governing lineage commitment. In spatial data, cortical-layer boundaries in the DLPFC remained sharp and continuous across all 8 sections after cross-slice alignment, outperforming STAligner and other spatial integration methods. The DLPFC cortical layers represent an anatomically defined, manually annotated gold standard for spatial integration evaluation, as each lamina contains distinct neuronal subtypes with well-characterized marker gene profiles that provide an objective ground truth against which the preservation of spatial coherence can be rigorously measured. In perturbation datasets, the integrated embedding of the Tahoe-100M lung-cancer subset retained drug-specific transcriptional contrasts sharp enough that drug perturbation correlation analysis recovered known target relationships and nominated candidate mechanisms of action for several drugs whose targets remain poorly characterized. For example, MEK inhibitors known to target the MAPK/ERK pathway showed strongly correlated transcriptional signatures in the integrated embedding, confirming that PRIME preserves the biological signal necessary to group mechanistically similar compounds. Compounds with less well-characterized targets showed strong correlations with known pathway signatures, providing testable mechanistic hypotheses that could guide future experimental validation. Taken together, these results are consistent with the central hypothesis that stabilizing local neighborhood structure during integration is essential for preserving the higher-order biological relationships, coherence of tissue architecture, and sharpness of perturbation responses.

Existing integration methods typically perform well on only one of these criteria. Classical MNN-based methods (MNN, fastMNN, and Scanorama) handle scRNA-seq effectively but ignore spatial coordinates. Recent spatial integration methods (GraphST and PRECAST) incorporate coordinates but lack adaptation to non-spatial data. Importantly, no batch effect correction method is specifically designed for perturbation screens, where the biological signal of interest is small relative to batch variation and is particularly vulnerable to overcorrection. PRIME instead treats expression-based consensus anchoring and coordinate-based spatial regularization as two graphs coupled within a single Laplacian objective, recovering each of these regimes as a special case: *β* = 0 reduces the framework to pure expression-based integration, increasing *α* emphasizes cross-slice alignment, and intermediate regimes smoothly interpolate. Classical MNN-based methods such as MNN and fastMNN are sensitive to any single projection’s idiosyncratic distortions, because projection noise translates directly into false anchors that corrupt the resulting correction. Harmony applies a global linear transformation to align batch centroids, which is effective when batch effects are approximately linear but can distort the local manifold topology underlying trajectory and clustering analyses. Deep generative approaches such as scVI avoid these assumptions through flexible neural network encoders, but require careful hyperparameter tuning, substantial computational resources, and are prone to instability when batch sizes are highly imbalanced. PRIME addresses these failure modes through a design that is simultaneously more robust than single-embedding matching, more topology-preserving than global linear correction, and more computationally efficient than neural network-based approaches.

Although PRIME achieved superior performance across scenarios, there are still several limitations warrant discussion. First, like all MNN-based methods, PRIME assumes at least partial overlap in cell states between batches; when integrated datasets share almost no common populations, consensus anchors become sparse and the method approaches the behavior of the underlying single-projection MNN. Second, the Laplacian regularization is intrinsically linear in the corrected embedding, and strongly nonlinear distortions such as cross-modality integration of RNA and ATAC-seq or protein data may still require deep learning-based approaches despite their higher computational cost^22^. Third, PRIME models only the batch covariates that are supplied as labels, thus latent sources of technical variation not captured by the user-provided batch annotation, such as ambient RNA contamination gradients or subtle protocol variations, are not explicitly inferred. Finally, the random-projection ensemble introduces stochasticity that users should control by fixing random seeds for reproducible outputs, beyond that, those very small ensembles (*K* < 3) could reduce the stability of consensus voting.

These limitations also point toward natural extensions. First, the framework can be adapted to streaming or online integration^41^, in which newly acquired batches are projected onto an existing consensus-anchor graph without recomputing the full ensemble from scratch, enabling incremental reference mapping against atlases that continuously grow. This online integration mode would also support a transfer-learning workflow in which newly profiled cells are mapped onto an existing PRIME-corrected reference, analogous to reference mapping approaches in which query datasets are projected onto a pre-built atlas embedding without perturbing the reference structure. Second, additional graphs can be incorporated into the Laplacian objective. For example, a lineage graph derived from trajectory priors or a perturbation-similarity graph for drug-target inference could give users principled control over the relative weight of different biological constraints^15^.

## Methods

### PRIME Algorithm

Given the normalized single cell expression matrix **X** ∈ ℝ^*n*×*d*^, PRIME generates *K* independent random projections^26,28^ into a lower-dimensional subspace 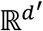 (default *d*′ = 128, *K* = 5). For each projection *k* ∈ {1, …, *K*}, a random matrix 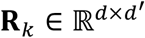 is drawn with entries sampled independently from *𝒩*(0,1/*d*′) ^2^, and the projected embedding is computed as:

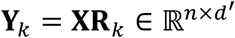

where **X** is the input normalized gene expression matrix, **R**_*k*_ is the *k*-th random projection matrix with Gaussian-distributed entries, **Y**_*k*_ is the *k*-th projected embedding, *K* is the total number of random projections, and *d*′ is the dimensionality of each projected space.

Random projection is motivated by the Johnson-Lindenstrauss lemma^26^, which guarantees that *n* points in high-dimensional Euclidean space can be embedded into a space of dimension *O*(𝜖^−2^log*n*) such that all pairwise distances are preserved up to a multiplicative factor of (1 ± 𝜖) with high probability. Each individual projection **Y**_*k*_ therefore provides a faithful but slightly distorted view of cell-cell geometry, and different random projections distort different regions of the manifold. Aggregating information across *K* independent projections averages out idiosyncratic distortions and yields more stable nearest-neighbor estimates than any single embedding.

### MNN anchor identification within each projection

Within each projection **Y**_*k*_, mutual nearest neighbors are identified between every pair of batches. For batches *a* and *b* with cell-index sets 𝔄 and 𝔅, and kNN_*k*_(⋅) denoting the nearest-neighbor operator in projection **Y**_*k*_, the MNN^18,20^ set is defined as:

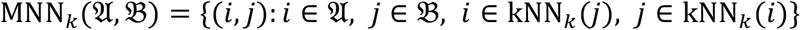

with default neighborhood size *k* = 20. The symmetric MNN condition requires each pair (*i, j*) to be a reciprocal nearest-neighbor correspondence: both cells find each other among their top-*k* neighbors. This criterion is more stringent than one-directional nearest-neighbor matching and reduces spurious anchors caused by sampling imbalance between batches.

### Ensemble consensus anchor selection

Single-projection MNN is known to produce false anchors when the projection distorts specific regions of the manifold; pairs that recur across independent random projections are geometrically robust correspondences. This is analogous to bootstrap aggregation^42^: individual estimators (single-projection MNNs) are noisy but unbiased, and averaging across *K* of them reduces variance without introducing systematic bias.

For every candidate pair (*i, j*), PRIME counts the number of projections in which the pair is supported as an MNN:

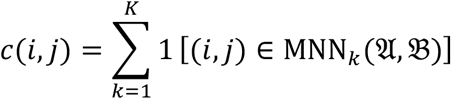

Pairs satisfying *c*(*i, j*) ≥ *τK* are retained as consensus anchors, where *τ* ∈ (0,1] is the support threshold (default *τ* = 0.5, i.e., majority voting). Let 𝒜(𝔄, 𝔅) denote the resulting consensus anchor set.

### Anchor-guided correction

For each consensus anchor pair (*i, j*) ∈ 𝒜(𝔄, 𝔅), a correction vector is computed in the original normalized expression space as:

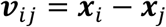

where *x*_*i*_, *x*_j_ ∈ ℝ^*d*^ are the gene expression profile of cells *i* and *j*. For each cell *c* in batch 𝔅, a smoothed correction vector is obtained as a Gaussian-weighted average over nearby anchors:

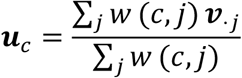

with weights *w*(*c, j*) = exp(−∥ ***x***_*c*_ − ***x***_j_ ∥^2^/(2*σ*^2^)Y; the bandwidth *σ* is set adaptively as the median distance from cell *c* to its 20 nearest anchors. The corrected embedding for cell *c* is:

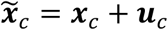

When more than two batches are present, the anchor graph is traversed in a greedy order determined by the size of the consensus anchor set between pairs: the pair with the largest |𝒜(𝔄, 𝔅)| is processed first, and each subsequently added batch is corrected against the running integrated reference. The final output is the corrected embedding 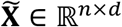, used for all downstream analyses.

### PRIME for Spatial Transcriptomics Data Integration

Spatial transcriptomics data resolves the physical position of each spot on a tissue section, which is absent from dissociated scRNA-seq. PRIME extends the core algorithm by constructing a spatial neighborhood graph and fusing it with the expression-based consensus graph through a graph-Laplacian spectral embedding^12,27^.

### Spatial neighborhood graph

Eigenvectors of the graph Laplacian with small eigenvalues encode the smoothest functions over the graph: cells connected by strong edges — whether because they are transcriptionally similar, spatially adjacent, or both — are mapped to nearby coordinates in **Z**. By combining **W**^*E*^ and **W**^*S*^ before constructing L, the resulting embedding respects both modalities simultaneously, avoiding both the spatial-blindness of expression-only integration and the expression-blindness of spatial-only smoothing.

For each tissue section, a *k*-nearest-neighbor graph is constructed from the two-dimensional spatial coordinates ***s***_*i*_ ∈ ℝ^2^ of each spot. With *k*_*s*_ spatial neighbors per spot (default *k*_*s*_ = 6, matching the hexagonal Visium lattice), the spatial similarity matrix is:

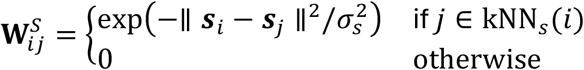

where *σ*_*s*_ is the median spatial distance across all kNN edges. **W**^*S*^ is symmetrized as **W**^*S*^ ← (**W**^*S*^ + (**W**^*S*^)^T^)/2.

### Expression-based consensus graph

From PRIME, for each pair of adjacent sections the consensus anchor set 𝔄 defines a binary expression-similarity matrix **W**^*E*^, where 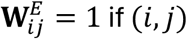 is a consensus anchor and 0 otherwise. Within a single section, **W**^*E*^ is populated from the ensemble kNN graph by thresholding *c*(*i, j*) ≥ *τK* on all within-section pairs, yielding a data-driven similarity graph that reflects transcriptomic proximity robust to projection noise.

### Graph-Laplacian fusion

The fused similarity graph combines expression and spatial similarity through a convex combination:

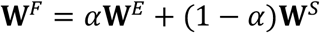

where *α* ∈ [0,1] controls the relative weight of expression versus spatial similarity (default *α* = 0.5; sensitivity analysis in Fig. S). The symmetric normalized Laplacian^27^ is:

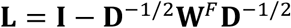

where **D** is the degree matrix with 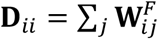 and I is the identity matrix.

### Spectral embedding

The PRIME integrated embedding **Z** ∈ ℝ^*n*×*p*^ is obtained by solving the eigenvalue problem:

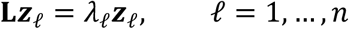

and stacking the eigenvectors **z**_2_, …, **z**_*p*+1_ associated with the *p* smallest non-zero eigenvalues (default *p* = 30) as columns of **Z**. The trivial eigenvector **z**_1_ (constant, *λ*_1_ = 0) is discarded.

### Datasets

We evaluated PRIME on five publicly available benchmarks spanning controlled cell-line mixtures, multi-study tumor atlases, developmental single-cell data, spatial transcriptomics, and large-scale perturbation profiling. Following common reporting practice in integration benchmark studies, we describe for each benchmark the original source, biological system, assay platform, batch structure, and any subset used in this work.

#### Controlled cell-line benchmark

As a controlled proof-of-concept benchmark, we used the official 10x Genomics Universal 3’ Jurkat, 293T, and 50:50 Jurkat:293T mixture datasets^31^. These libraries contain approximately 3,258 Jurkat cells, 2,885 293T cells, and 3,400 cells in the mixed library, respectively. We treated the two pure cell-line libraries and the 1:1 mixture library as three batches with known ground-truth identities, providing a simple setting in which successful integration should remove batch structure while preserving the separation between the two cell lines.

#### Human lung cancer compendium

For large-scale single-cell benchmarking in a clinically heterogeneous setting, we used a curated human lung cancer scRNA-seq compendium^43^ assembled from multiple publicly available studies comprising malignant, immune, and stromal populations. In the main text, we summarize the merged benchmark by its total number of cells, donors, and studies after harmonization and quality control.

#### Human hematopoiesis benchmark

To evaluate preservation of developmental structure after integration, we used a human bone marrow single-cell RNA-seq dataset spanning healthy donors and covering hematopoietic stem/progenitor cells together with lineage-committed descendants^44^. For trajectory-focused analyses, we further restricted the dataset to the hematopoietic compartments relevant to the erythro-myeloid continuum.

#### DLPFC spatial transcriptomics benchmark

For spatial integration, we used the human dorsolateral prefrontal cortex (DLPFC) Visium dataset originally described by Maynard et al.^13^ This resource contains 12 tissue sections from 3 neurotypical adult donors, with two pairs of spatially adjacent serial sections per donor, and includes manually curated cortical layer labels that provide a biologically grounded reference for evaluating spatial integration. Processed spot-level data and layer annotations were obtained through the spatialLIBD resource^45^. Because each section spans the cortical laminae and white matter, this dataset provides a stringent benchmark for assessing both cross-section alignment and preservation of within-section spatial coherence.

#### Tahoe-100M perturbation benchmark

To assess atlas-scale perturbation integration, we used Tahoe-100M, a public single-cell perturbation atlas comprising more than 100 million transcriptomic profiles from 50 cancer cell lines exposed to approximately 1,100 small-molecule perturbations^17^. The dataset was accessed through the Arc Virtual Cell Atlas and the public Hugging Face release, which provide raw expression data together with sample-, drug-, cell-line-, and observation-level metadata. Because the full resource is substantially larger than is practical for routine benchmarking, we analyzed a lung-cancer-focused subset containing approximately 5 million cells across selected drug treatments and matched DMSO controls.

#### Common preprocessing and metadata handling

For all datasets, we started from publicly available count matrices and accompanying metadata. Quality control, normalization, highly variable gene selection, and dimensionality reduction were then performed using a unified preprocessing workflow described below, so that performance differences reflected the integration method rather than dataset-specific preprocessing choices.

### Benchmark metrics

We evaluated integration performance with a panel of metrics grouped into three categories: (i) batch correction, (ii) biological conservation, and (iii) for spatial data, spatial conservation. All scIB-family metrics were computed using the scib-metrics Python package following the protocol of Luecken et al^5^. Each metric is scaled to [0,1], with higher values indicating better integration.

#### Batch correction metrics

These metrics quantify the extent to which batch-associated variation has been removed from the integrated embedding.

##### Graph iLISI^20^ (integration Local Inverse Simpson’s Index)

Measures, at the level of each cell’s local neighborhood in the kNN graph, the effective number of batches represented. For cell *i* with neighborhood-level batch distribution *p* = (*p*_1_, …, *p*_𝔅_) over 𝔅 batches, the local iLISI is 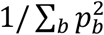. An ideally integrated embedding in which every neighborhood contains all batches in equal proportion yields iLISI å 𝔅; complete batch separation yields iLISI ≈ 1. Values are rescaled to [0,1] across cells.

##### kBET^4C^ (k-nearest-neighbor batch effect test)

Performs a *χ*^2^ test at each cell’s kNN neighborhood, asking whether the batch composition of the neighborhood differs significantly from the global batch composition. The reported score is 1 minus the rejection rate across cells, so well-mixed embeddings (few rejections) score close to 1.

##### Graph connectivity^5^

For each cell type, quantifies whether cells of that type form a single connected component in the kNN graph of the integrated embedding. It is defined as the mean, across cell types, of the largest-connected-component fraction within each cell type. A score of 1 indicates that every cell-type-restricted subgraph is fully connected — the expected behavior when batch effects have been removed within each cell type.

##### Batch ASW^47^ (average silhouette width)

Computes the silhouette coefficient of each cell with respect to batch labels, then inverts and rescales so that low silhouette magnitude (weak batch-based separation) yields a higher score.

##### Principal component regression^4C^ (PCR)

Regresses the top principal components of the integrated embedding against the batch variable and reports 1 − ℝ^2^ weighted by the explained variance of each PC. Good integration reduces the fraction of variance explainable by batch.

#### Biological conservation metrics

These metrics assess how well the integrated embedding preserves biological structure — cell identity, lineage relationships, and gene programs.

##### NMI (normalized mutual information)

Mutual information between Leiden clusters computed on the integrated embedding and ground-truth cell-type labels, normalized to [0,1]. We report the maximum NMI over a sweep of Leiden resolutions to eliminate resolution dependence.

##### ARI (adjusted Rand index)

Measures the agreement between Leiden clustering and ground-truth labels, corrected for chance agreement. Unlike NMI, ARI does not inflate with cluster number, providing a complementary view.

##### Cell-type ASW

Silhouette coefficient of each cell with respect to its cell-type label; higher values indicate that cells cluster tightly with others of the same type and are well separated from other types.

##### Graph cLISI (cell-type Local Inverse Simpson’s Index)

The cell-type analog of iLISI: for each cell’s kNN neighborhood, it computes the effective number of distinct cell-type labels, rescaled so that neighborhoods dominated by a single cell type (good cell-identity preservation) score close to 1.

##### Isolated-label F1 and ASW

Evaluates whether rare cell types — specifically, those present in only a small number of batches — remain distinguishable after integration. This metric explicitly penalizes overcorrection that merges rare populations into larger, batch-ubiquitous clusters.

##### XLC

To evaluate whether an integrated embedding preserves the expected laminar continuity of spatial transcriptomics data, we designed a cross-layer-neighborhood continuity (XLC) score. Unlike conventional clustering metrics such as ARI or NMI, which treat all annotation mismatches equally, XLC explicitly uses the ordered structure of cortical layers. For each spot *i*, we identified its *k*-nearest neighbors 𝔑_*k*_(*i*) in the integrated embedding space. When evaluating cross-slice integration, neighbors were restricted to spots from different slices. The local ordinal continuity score for spot *i* was defined as

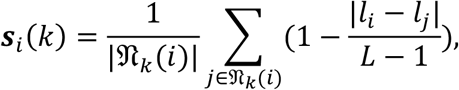

where *l*_*i*_ and *l*_j_ denote the ordinal layer ranks of spots *i* and *j*, respectively, and *L* = 7 is the total number of ordered laminar categories. This score equals 1 when neighboring spots have identical layer annotations and decreases linearly as their layer-rank distance increases. To avoid dominance by layers with more spots, we computed a macro-averaged score across layers:

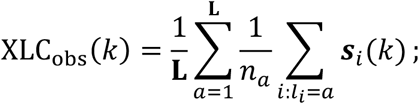

where *n*_*a*_ is the number of spots assigned to layer rank *a*. Higher XLC values indicate that nearby spots in the integrated embedding tend to have adjacent laminar annotations, suggesting better preservation of biologically meaningful spatial continuity across slices.

### Benchmark framework

#### Preprocessing

Raw count matrices from each batch are normalized by library-size scaling followed by log(1 + *x*) transformation. Highly variable genes (HVGs) are selected jointly across batches using the `Scanpy flavor=“seurat_v3”` procedure, retaining the top 2,000 genes by dispersion (value chosen per dataset; see Table S1). The concatenated HVG matrix 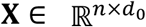 (with *n* cells and *d*_0_ HVGs) is reduced by principal component analysis (PCA) to *d* principal components (default *d* = 50), yielding the input matrix **X** ∈ ℝ^*n*×*d*^ used in all subsequent steps.

#### Benchmarking methods

To comprehensively evaluate the performance of PRIME, we benchmarked against ten methods for scRNA-seq data integration and five methods for spatial transcriptomics integration, covering MNN/anchor-based, graph-based, matrix-factorization, and deep-generative. We used the default parameter settings for all methods for data integration.

#### Trajectory inference

To evaluate the capability of estimating the underlying trajectory of the methods, we conducted trajectory inference of a human hematopoiesis dataset. We applied Monocle3^36^, which directly infers the trajectory based on the UMAP embeddings generated by the integrated embeddings, and the root has been manually assigned to the Hematopoietic Stem and Progenitor Cells (HSPCs).

## Data Availability

There is no new data generated in this study. All datasets used for benchmarking are publicly available and described in the Methods section.

## Code Availability

The code for PRIME algorithm and spatial extension can be available at https://github.com/wan-mlab/PRIME. And the code for all the benchmarking analyses, including data analysis, metric computation, and figure preparation, is also included in the same repository.

## Acknowledgements

Research reported in this publication was supported by the U.S. National Science Foundation under Award Number 2500836, and the Office Of The Director, National Institutes Of Health of the National Institutes of Health under Award Number R03OD038391. This work was also partially supported by the National Institute of General Medical Sciences of the National Institutes of Health under Award Numbers P20GM103427. This study was in part financially supported by the Child Health Research Institute at UNMC/Children’s Nebraska. This research was supported by the State of Nebraska through the Pediatric Cancer Research Group, part of the Child Health Research Institute. The content is solely the responsibility of the authors and does not necessarily represent the official views of the funding organizations.

## Author Contribution

W.S. and W.-X.C. conceived and designed the study. W.-X.C. developed the methods. W.-X.C. and W.-X.F. performed the experiments and analyzed the data. W.S. and W.-X.C. participated in writing the manuscript. W.S., W.-X.C., W.-X.F. and W.J. revised the manuscript. The manuscript was approved by all authors.

## References

1. Rood, J. E. et al. The Human Cell Atlas: from a cell census to a unified foundation model. Nature 637, 1065–1071 (2025).

2. Method of the Year 2020: spatially resolved transcriptomics. Nat Methods 18, 1–1 (2021).

3. Tabula Sapiens Consortium. The Tabula Sapiens: a multiple-organ, single-cell transcriptomic atlas of humans. Science 376, eabl4896 (2022).

4. Regev, A. et al. The Human Cell Atlas. eLife 6, e27041 (2017).

5. Luecken, M. D. et al. Benchmarking atlas-level data integration in single-cell genomics. Nat Methods 19, 41–50 (2022).

6. Tran, H. T. N. et al. A benchmark of batch-effect correction methods for single-cell RNA sequencing data. Genome Biol 21, 12 (2020).

7. Wang, H., Leskovec, J. & Regev, A. Limitations of cell embedding metrics assessed using drifting islands. Nature Biotechnology 44, 574–577 (2025).

8. Trapnell, C. et al. The dynamics and regulators of cell fate decisions are revealed by pseudotemporal ordering of single cells. Nature Biotechnology 32, 381–386 (2014).

9. Qiu, X. et al. Reversed graph embedding resolves complex single-cell trajectories. Nature Methods 14, 979–982 (2017).

10. Wolf, F. A. et al. PAGA: graph abstraction reconciles clustering with trajectory inference through a topology preserving map of single cells. Genome Biology 20, 59 (2019).

11. Saelens, W., Cannoodt, R., Todorov, H. & Saeys, Y. A comparison of single-cell trajectory inference methods. Nat Biotechnol 37, 547–554 (2019).

12. Ståhl, P. L. et al. Visualization and analysis of gene expression in tissue sections by spatial transcriptomics. Science 353, 78–82 (2016).

13. Maynard, K. R. et al. Transcriptome-scale spatial gene expression in the human dorsolateral prefrontal cortex. Nature Neuroscience 24, 425–436 (2021).

14. Rao, A., Barkley, D., França, G. S. & Yanai, I. Exploring tissue architecture using spatial transcriptomics. Nature 596, 211–220 (2021).

15. Dixit, A. et al. Perturb-Seq: Dissecting molecular circuits with scalable single-cell RNA profiling of pooled genetic screens. Cell 167, 1853–1866 (2016).

16. Subramanian, A. et al. A next generation connectivity map: L1000 platform and the first 1,000,000 profiles. Cell 171, 1437–1452 (2017).

17. Zhang, J. et al. Tahoe-100M: a giga-scale single-cell perturbation atlas for context-dependent gene function and cellular modeling. bioRxiv https://doi.org/10.1101/2025.02.20.639398 (2025) doi:10.1101/2025.02.20.639398.

18. Haghverdi, L., Lun, A. T. L., Morgan, M. D. & Marioni, J. C. Batch effects in single-cell RNA-sequencing data are corrected by matching mutual nearest neighbors. Nat Biotechnol 36, 421–427 (2018).

19. Butler, A., Hoffman, P., Smibert, P., Papalexi, E. & Satija, R. Integrating single-cell transcriptomic data across different conditions, technologies, and species. Nat Biotechnol 36, 411–420 (2018).

20. Korsunsky, I. et al. Fast, sensitive and accurate integration of single-cell data with Harmony. Nat Methods 16, 1289–1296 (2019).

21. Polański, K. et al. BBKNN: fast batch alignment of single cell transcriptomes. Bioinformatics 36, 964–965 (2020).

22. Lopez, R., Regier, J., Cole, M. B., Jordan, M. I. & Yosef, N. Deep generative modeling for single-cell transcriptomics. Nat Methods 15, 1053–1058 (2018).

23. Liu, X., Zeira, R. & Raphael, B. J. Partial alignment of multi-slice spatially resolved transcriptomics data. Genome Research 33, 1124–1132 (2023).

24. Long, Y. et al. Spatially informed clustering, integration, and deconvolution of spatial transcriptomics with GraphST. Nat Commun 14, 1155 (2023).

25. Clifton, K. et al. STalign: Alignment of spatial transcriptomics data using diffeomorphic metric mapping. Nat Commun 14, 8123 (2023).

26. Conference in Modern Analysis and Probability (1982 : Yale University). Conference in Modern Analysis and Probability. (Providence, R.I. : American Mathematical Society, 1984).

27. Belkin, M. & Niyogi, P. Laplacian Eigenmaps for Dimensionality Reduction and Data Representation. Neural Computation 15, 1373–1396 (2003).

28. Achlioptas, D. Database-friendly random projections: Johnson-Lindenstrauss with binary coins. Journal of Computer and System Sciences 66, 671–687 (2003).

29. Wan, S., Kim, J. & Won, K. J. SHARP: hyperfast and accurate processing of single-cell RNA-seq data via ensemble random projection. Genome Res 30, 205–213 (2020).

30. Zeira, R., Land, M., Strzalkowski, A. & Raphael, B. J. Alignment and integration of spatial transcriptomics data. Nat Methods 19, 567–575 (2022).

31. 50%:50% Jurkat:293T Cell Mixture. 10x Genomics https://www.10xgenomics.com/datasets/50-percent-50-percent-jurkat-293-t-cell-mixture-1-standard-1-1-0.

32. Lambrechts, D. et al. Phenotype molding of stromal cells in the lung tumor microenvironment. Nat Med 24, 1277–1289 (2018).

33. Hie, B., Bryson, B. & Berger, B. Efficient integration of heterogeneous single-cell transcriptomes using Scanorama. Nat Biotechnol 37, 685–691 (2019).

34. Johnson, W. E., Li, C. & Rabinovic, A. Adjusting batch effects in microarray expression data using empirical Bayes methods. Biostatistics 8, 118–127 (2007).

35. Stuart, T. et al. Comprehensive Integration of Single-Cell Data. Cell 177, 1888-1902.e21 (2019).

36. Cao, J. et al. The single-cell transcriptional landscape of mammalian organogenesis. Nature 566, 496–502 (2019).

37. Velten, L. et al. Human haematopoietic stem cell lineage commitment is a continuous process. Nat Cell Biol 19, 271–281 (2017).

38. Pellin, D. et al. A comprehensive single cell transcriptional landscape of human hematopoietic progenitors. Nat Commun 10, 2395 (2019).

39. Maynard, K. R. et al. Transcriptome-scale spatial gene expression in the human dorsolateral prefrontal cortex. Nat Neurosci 24, 425–436 (2021).

40. Liu, W. et al. Probabilistic embedding, clustering, and alignment for integrating spatial transcriptomics data with PRECAST. Nat Commun 14, 296 (2023).

41. Lotfollahi, M. et al. Mapping single-cell data to reference atlases by transfer learning. Nat Biotechnol 40, 121–130 (2022).

42. Breiman, L. Bagging predictors. Mach Learn 24, 123–140 (1996).

43. Sikkema, L. et al. An integrated cell atlas of the lung in health and disease. Nat Med 29, 1563–1577 (2023).

44. A single cell immune cell atlas of human hematopoietic system - Overview - HCA Data Explorer. https://explore.data.humancellatlas.org/projects/cc95ff89-2e68-4a08-a234-480eca21ce79.

45. Pardo, B. et al. spatialLIBD: an R/Bioconductor package to visualize spatially-resolved transcriptomics data. BMC Genomics 23, 434 (2022).

46. Büttner, M., Miao, Z., Wolf, F. A., Teichmann, S. A. & Theis, F. J. A test metric for assessing single-cell RNA-seq batch correction. Nat Methods 16, 43–49 (2019).

47. Rousseeuw, P. J. Silhouettes: A graphical aid to the interpretation and validation of cluster analysis. Journal of Computational and Applied Mathematics 20, 53–65 (1987).

