## Supplementary figures and images for "Atlas-Level Single-Cell and Spatial Transcriptomics Data Integration via PRIME"

### Supplementary Figure 1

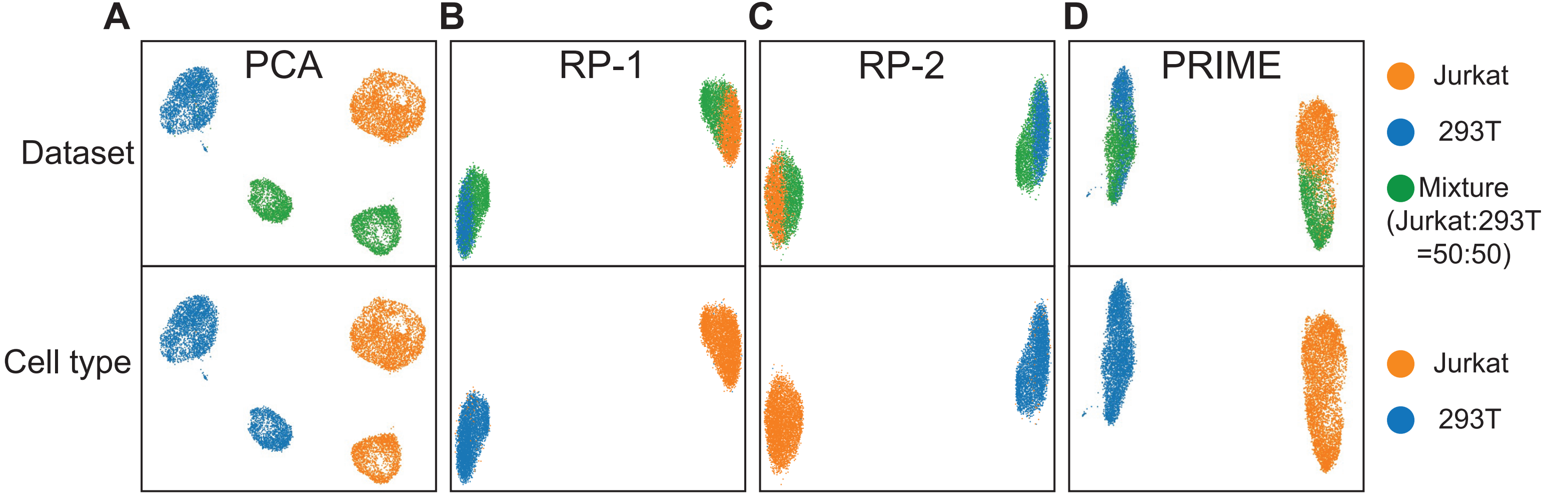

### Supplementary Figure 2

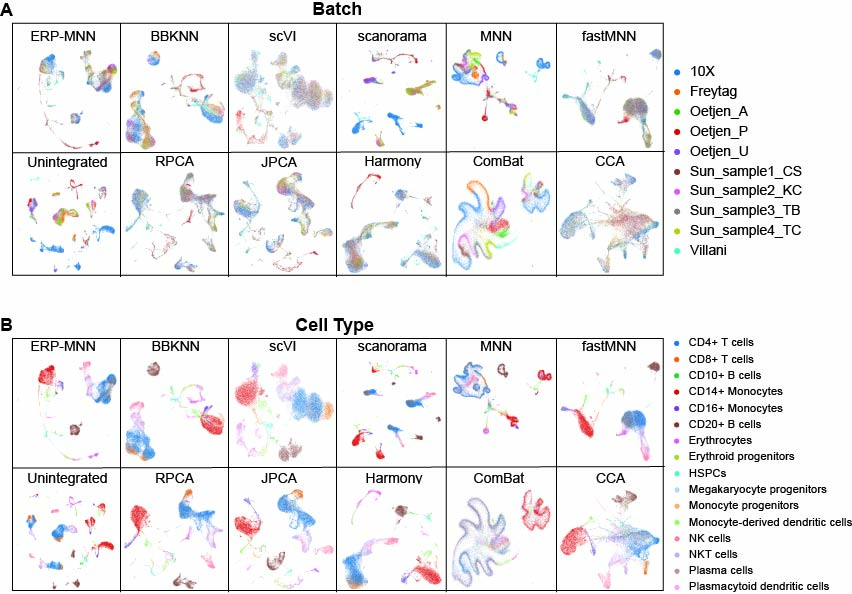
